# A novel signature derived from immunoregulatory and hypoxia genes predicts prognosis in liver and five other cancers

**DOI:** 10.1101/442921

**Authors:** Wai Hoong Chang, Donall Forde, Alvina G. Lai

## Abstract

**Background:** Despite much progress in cancer research, it’s incidence and mortality continue to rise. A robust biomarker that would predict tumor behavior is highly desirable and could improve patient treatment and prognosis.

**Methods:** In a retrospective bioinformatics analysis involving patients with liver cancer (n=839), we developed a prognostic signature consisting of 45 genes associated with tumor-infiltrating lymphocytes and cellular responses to hypoxia. From this gene set, we were able to identify a second prognostic signature comprised of 8 genes. Its performance was further validated in five other cancers: head and neck (n=520), renal papillary cell (n=290), lung (n=515), pancreas (n=178) and endometrial (n=370).

**Findings:** The 45-gene signature predicted overall survival in three liver cancer cohorts: hazard ratio (HR)=1.82, P=0.006; HR=1.84, P=0.008 and HR=2.67, P=0.003. Additionally, the reduced 8-gene signature was sufficient and effective in predicting survival in liver and five other cancers: liver (HR=2.36, P=0.0003; HR=2.43, P=0.0002 and HR=3.45, P=0.0007), head and neck (HR=1.64, P=0.004), renal papillary cell (HR=2.31, P=0.04), lung (HR=1.45, P=0.03), pancreas (HR=1.96, P=0.006) and endometrial (HR=2.33, P=0.003). Receiver operating characteristic analyses demonstrated both signature’s superior performance over current tumor staging parameters. Multivariate Cox regression analyses revealed that both 45-gene and 8-gene signatures were independent of other clinicopathological features in these cancers. Combining the gene signatures with somatic mutation profiles increased their prognostic ability.

**Conclusions:** This study, to our knowledge, is the first to identify a gene signature uniting both tumor hypoxia and lymphocytic infiltration as a prognostic determinant in six cancer types (n=2,712). The 8-gene signature can be used for patient risk stratification by incorporating hypoxia information to aid clinical decision making.

## Background

Hepatocellular carcinoma (HCC) is the sixth most common cancer and the second leading cause of cancer-related mortality worldwide[1]. Due to an initially asymptomatic disease course, this aggressive cancer once diagnosed has especially poor outcomes. The etiological risk factors for HCC varies across geographical locations[2]. This pattern mirrors the burden of viral hepatitis in Asia and Africa. Whereas in the Western world, the risk can be attributed more to alcoholic and non-alcoholic steatohepatitis. Therapy for HCC, involving either surgery or radiological ablation, is most effective when the cancer is detected early. Prompt diagnosis, however, requires regular liver imaging, usually six-monthly ultrasound scans, which is both resource intensive and dependent upon identification of otherwise silent risk factors. Once diagnosed, curative liver transplantation is based on tumor size and restricted to patients where the primary cancer is thought unlikely to recur in the new liver. The use of size as the only prognostic criterion precludes therapy in patients with large, indolent cancers that have a low probability of recurrence and many centers are questioning whether additional data on tumor biology would add value to the diagnostic pathway.

Gene signatures obtained from tumor that are derived from common oncogenic pathways can be used to risk-stratify patients to provide individualized care. Despite genomic instability driving intertumoral heterogeneity, solid malignancies do share two common characteristics. Solid tumors are often hypoxic due to aberrant vasculature resulting in metabolic shifts towards aerobic glycolysis known as the Warburg effect[3,4]. Tumor hypoxia is associated with metastases and aggressive phenotypes that influence clinical outcomes[5,6]. Additionally, within tumor cells one can find lymphocytes termed tumor-infiltrating lymphocytes. These can affect patient prognosis in multiple cancers[7,8]. A subset of lymphocytic infiltrates known as the FoxP3^+^ regulatory T cells (Tregs) function to suppress cytotoxic T cells activity to maintain tumor tolerance. Increases in Tregs contributes to unfavorable prognosis in multiple solid tumors[9]. Importantly, infiltrating Tregs within the tumor milieu can be affected by hypoxia where the latter facilitates Tregs recruitment to promote angiogenesis and maintain growth[10]. Another category of T cells known as the cytotoxic CD8^+^ T cells are required to kill cancer cells. But they can be made ineffective by Tregs or be rendered dysfunctional when exhausted[8]. Together, accumulation of Tregs and exhausted CD8^+^ T cells further exacerbate the immunosuppressive functions in solid tumors.

By taking advantage of the intricate link between hypoxia and Treg infiltration in solid malignancies, we systematically developed two prognostic gene signatures for risk stratification. Both signatures can accurately predict high-risk patients as confirmed by multi-cohort validations in six cancer types to support their validity and immediate clinical application.

## Results

### Hypoxia is associated with immunosuppression through enhanced expression of Tregs and CD8^+^ T cells genes

Since both hypoxia and high density of infiltrating Tregs are linked to poor clinical outcomes, we hypothesized that these features could be employed to predict prognosis in patients with hepatocellular carcinoma (HCC). Hypoxia inducible factors (HIFs) are transcription factors (TFs) that play key roles in regulating cellular responses to hypoxia[11]. The alpha subunits (HIF-1α and −2α) heterodimerize with the beta subunit (HIF-1β) to orchestrate physiological responses in hypoxia[12]. To develop our gene signature uniting hypoxia and tumor-infiltrating T-cells, we utilized three gene sets: 1) pan-cancer hypoxia genes (52 genes)[6]; 2) HCC-infiltrating Tregs (401 genes) and 3) HCC-infiltrating exhausted CD8^+^ T cells (82 genes)[8](Fig.1). To determine which of these genes were bound by HIFs, we used a HepG2 hepatoma HIF-1α/2α chromatin immunoprecipitation sequencing (ChIP-seq) dataset (GSE120885) (Fig.1)

**Figure 1.**
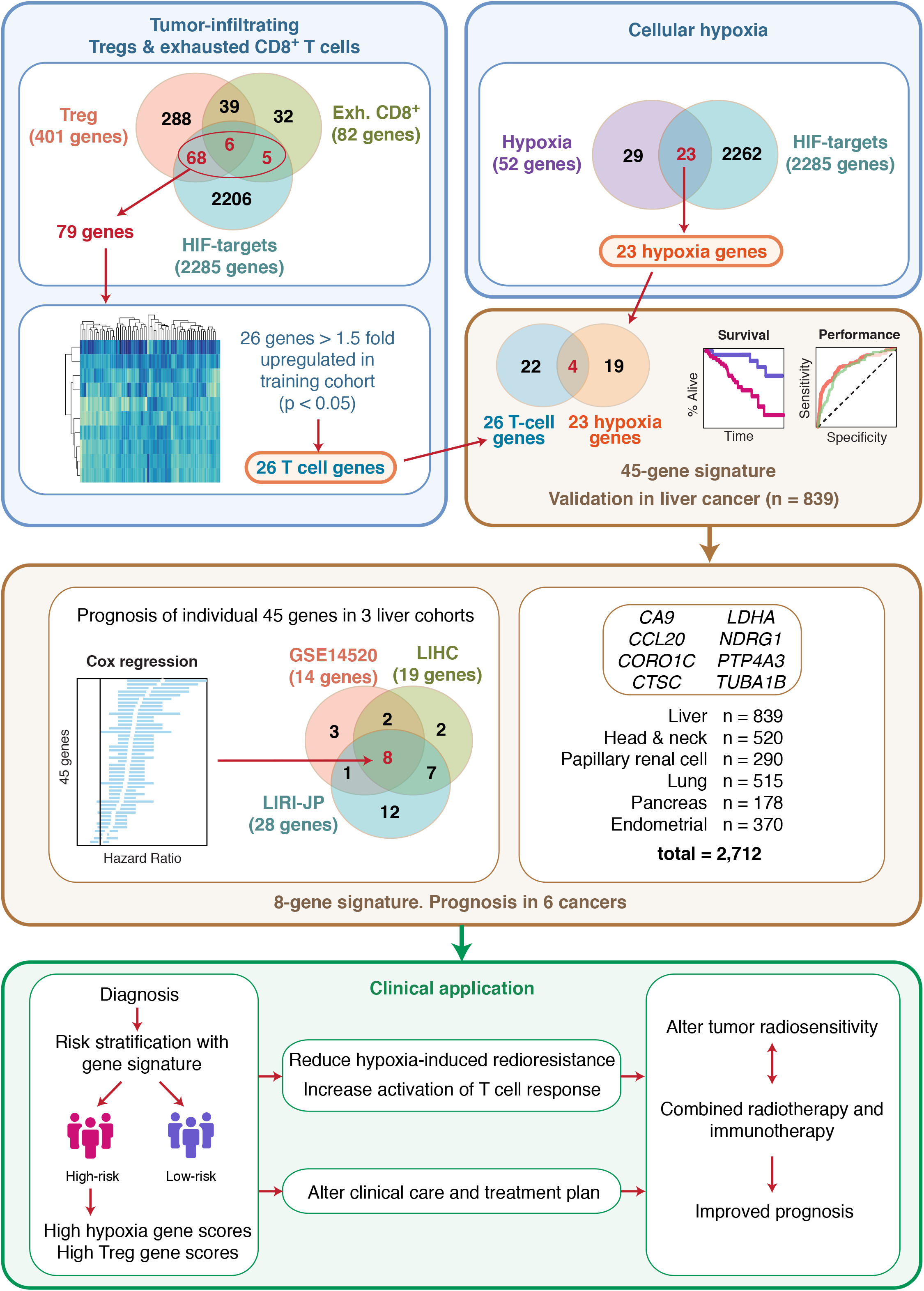
Schematic diagram of the study design and development of gene signatures. A liver cancer cohort (GSE14520) was used to define the first 45-gene signature. Briefly, 79 tumor-infiltrating T-cell genes were identified as HIF targets using a HIF-1α/2α ChIP-seq dataset. Of these 79 genes, 26 genes were > 1.5-fold upregulated in GSE14520. Independently, 23 hypoxia genes were identified as HIF targets. Uniting the 23 hypoxia-HIF genes and 26 T-cell-HIF genes resulted in 45 unique genes representing the first gene signature. Cox regression analyses of individual 45 genes in each of the three liver cancer cohorts (GSE14520, TCGA-LIHC and LIRI-JP) revealed a common prognostic set consisting of 8 genes that represent the second signature. This 8-gene signature is further validated in liver and five other cancers using Kaplan-Meier, Cox regression and receiver operating characteristic analyses.

To determine the extent of hypoxia in cancers, we interrogated *in vivo* mRNA expression patterns of the 52 hypoxia genes[6] in 25 cancer types including HCC (n=20,662) retrieved from The Cancer Genome Atlas (TCGA)[13](Table S1). Hypoxia scores for each tumor and non-tumor samples were determined by obtaining the mean expression values (log_2_) of the 52 hypoxia genes[6]. We observed that tumor samples were significantly more hypoxic than non-tumor samples in 20 out of 25 cancers, which included HCC (Fig.S1A). Multidimensional scaling analyses revealed that the 52 genes can distinguish tumor from non-tumor samples in these cancers, hence hypoxic transcriptional states can be used as a proxy for identifying cancerous cells (Fig.S1B).

We predict that patients with more hypoxic tumors would have higher expression of tumor-infiltrating T cell genes since hypoxia could promote the maintenance of immunological escape via the suppressive function of Tregs[10]. The HCC-infiltrating T-cell gene set uniting both Tregs and exhausted CD8^+^ T cells consisted of 438 genes collectively, with 45 genes found to be in common between the two cell types (Fig.1). Infiltrating T cell expression scores for each patient were calculated as mean expression values (log_2_) of the 438 genes (Fig.1). Patient hypoxia scores significantly correlated with tumor-infiltrating T cell scores in three independent HCC cohorts (n = 839; Table S2): GSE14520 (ρ=0.564, P<0.0001); TCGA-LIHC (ρ=0.390, P<0.0001) and LIRI-JP (ρ=0.633, P<0.0001) (Fig.S1C), suggesting that highly hypoxic tumors have transcriptional profiles associated with immunosuppressive functions.

Since HIFs are the key TFs that regulate numerous important aspects of oncogenesis[14], we hypothesize that identifying genes that are directly bound by HIFs may have more profound implications on prognosis. Using the HIF ChIP-seq dataset, we observed that 79 of the 438 tumor-infiltrating T cell genes were direct HIF targets (Fig.1). Notably, Spearman’s correlation coefficients between the 79 genes and hypoxia score were significantly higher: GSE14520 (ρ=0.669, P<0.0001), TCGA-LIHC (ρ=0.469, P<0.0001) and LIRI-JP (ρ=0.694, P<0.0001) (Fig.S1D), suggesting a role for HIFs in regulating tumor lymphocytic infiltration in HCC.

Levels of T cell infiltration were corrected by dividing the expression of each of the 79 genes with *CD3D* expression. Of the 79 genes, 26 were upregulated more than 1.5-fold in the training cohort, GSE14520 (Fig.S1E). On the other hand, ChIP-seq analysis revealed that 23 of the 52 hypoxia genes mentioned earlier were direct HIF targets (Fig.1). The final gene set was comprised of 45 genes that encompass 26 tumor-infiltrating lymphocyte genes and 23 hypoxia genes, with 4 genes being in common (Fig.1; Table S3).

### The 45-gene signature is strongly associated with poor overall survival in HCC

We next assessed the ability of the 45-gene signature (HCC45) to predict overall survival (OS) in three independent HCC cohorts (n=839). The signature can successfully discriminate between tumor and non-tumor samples in these cohorts (Fig.S2A). Tumor samples were notably less tightly clustered implying significant intertumoral heterogeneity (Fig.S2A). We next determined HCC45 scores, calculated for each patient as the mean expression of the 45 genes. Patients were ranked according to their HCC45 scores and divided into high- or low-risk groups using the median cutoff. The OS rates were significantly reduced in high-risk patient groups with log-rank P values of 0.012, 0.0042 and 0.0043 in the GSE14520, TCGA-LIHC and LIRI-JP cohorts respectively (Fig.2A). Disease-free survival rates were also significantly lower in high-risk patients (P=0.026) (Fig.S2B).

**Figure 2.**
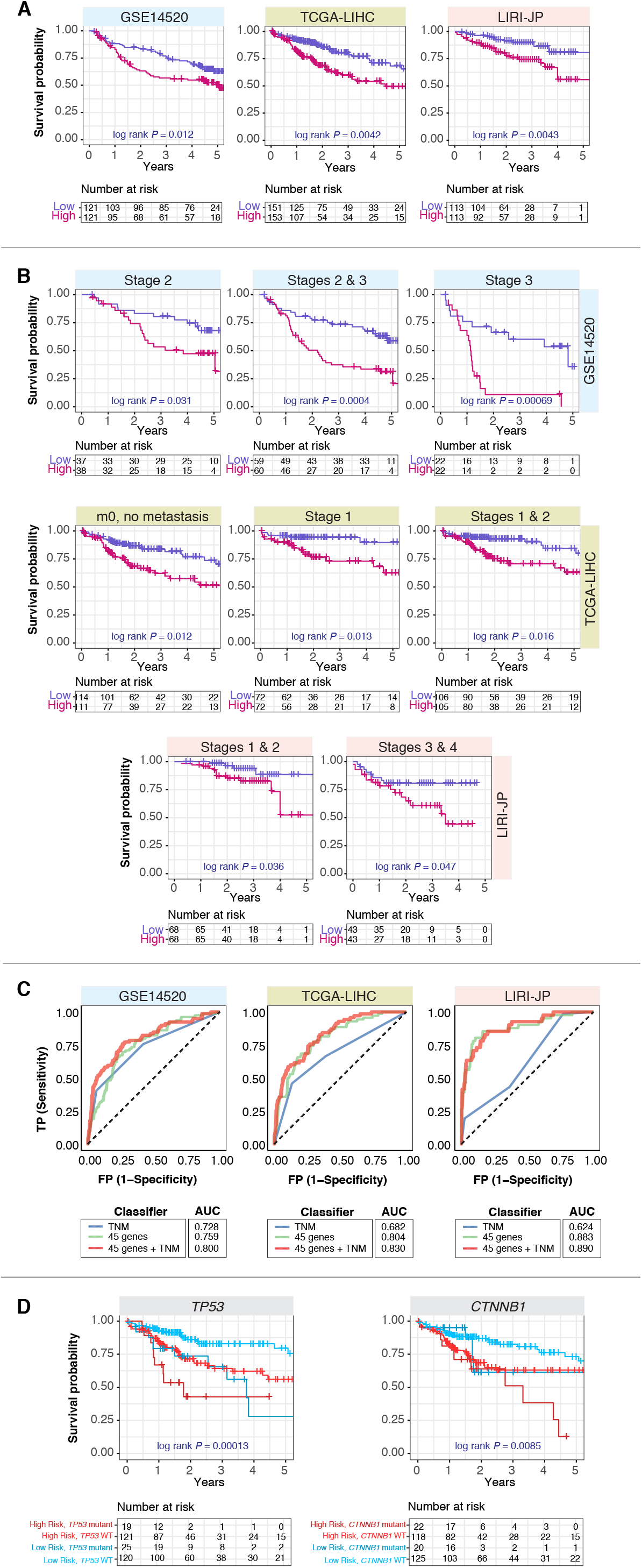
Patient stratification using the 45-gene signature in HCC cohorts. **(A)** Kaplan-Meier plots of overall survival in HCC patients across three cohorts stratified into low- and high-risk groups using the 45-gene signature. P-values were calculated from the log-rank test. **(B)** Kaplan-Meier plots show independence of the signature over current staging systems in HCC cohorts. Patients were sub-grouped according to TNM stages and further stratified using the 45-gene signature. The signature successfully identified high-risk patients in different TNM stages. P-values were calculated from the log-rank test. **(C)** Analysis of specificity and sensitivity of the signature in HCC cohorts with receiver operating characteristic (ROC). Plots depict comparison of ROC curves of signature and clinical tumor staging parameters. The signature demonstrates an incremental value over current staging systems. AUC: area under the curve. TNM: tumor, node, metastasis staging. **(D)** Kaplan-Meier plots depicting combined relation of *TP53* or *CTNNB1* mutation status with the signature on overall survival in HCC.

To test the independence of HCC45 over current staging systems, we performed subgroup analysis by applying the signature to patients within each tumor, node metastasis (TNM) stages. The signature successfully identified high-risk patients, particularly among early-stage (1 and/or 2) patients (Fig.2B). To evaluate the predictive performance of HCC45 on 5-year OS, we performed the receiver operating characteristic (ROC) analysis in comparison with tumor staging parameters. The area under the curves (AUC) for HCC45 were 0.759 (GSE14520), 0.804 (TCGA-LIHC) and 0.883 (LIRI-JP) (Fig.2C). All HCC45 AUCs were higher than that of TNM staging (Fig.2C). Moreover, combining HCC45 with TNM staging considerably improved its predictive ability: AUC = 0.800 (GSE14520), 0.830 (TCGA-LIHC) and 0.890 (LIRI-JP) (Fig.2C).

We assessed two common mutations in HCC, *TP53* and *CTNNB1*. Mutation in the tumor suppressor *TP53* is associated with poor prognosis[15,16]. Results on β-catenin *CTNNB1* mutations have been mixed; some demonstrating poor outcomes[17,18], while others favorable prognosis[19]. Both *TP53* (P<0.001) and *CTNNB1* (P=0.025) mutations were associated with poor prognosis (Table 1). Mortality in patients with high HCC45 scores and *TP53* mutation were ~45% higher than in patients with low HCC45 scores harboring wild-type *TP53* at 2 years (Fig.2D). Likewise, joint relation of HCC45 and *CTNNB1* predicted a ~35% increase in mortality at 4 years between the lowest and highest risk groups (Fig.2D).

**Table 1.** Univariate and multivariate Cox proportional hazard regression analysis of risk factors associated with overall survival.

|  | Hazard Ratio (95% CI) | P-value |
| --- | --- | --- |
| **GSE14520** |  |  |
|  | **Univariate** |  |
| 45-gene signature (high vs. low risk) | 1.823 (1.18 - 2.816) | **0.00683** |
| 8-gene signature (high vs. low risk) | 2.36 (1.49 - 3.74) | **0.00026** |
| Tumour size (> 5cm vs. < 5cm) | 2.159 (1.406 - 3.316) | **0.00044** |
| Cirrhosis (yes vs. no) | 4.665 (1.147 - 18.97) | **0.0314** |
| TNM staging (II & III vs. I) | 2.26 (1.706 - 2.994) | **1.30E-08** |
| BCLC staging (B & C vs. A & 0) | 2.181 (1.722 - 2.762) | **9.86E-11** |
| AFP (> 300ng/mL vs. < 300ng/mL) | 1.606 (1.049 - 2.46) | **0.0293** |
|  | **Multivariate (45-gene signature)** |  |
| 45-gene signature (high vs. low risk) | 1.8864 (1.2088 - 2.944) | **0.00519** |
| Tumour size (> 5cm vs. < 5cm) | 0.9487 (0.5605 - 1.606) | 0.8445 |
| Cirrhosis (yes vs. no) | 4.5927 (1.1084 - 19.031) | **0.03557** |
| TNM staging (II & III vs. I) | 1.4359 (1.2752 - 2.114) | **0.04684** |
| BCLC staging (B & C vs. A & 0) | 1.7808 (1.2521 - 2.533) | **0.00132** |
| AFP (> 300ng/mL vs. < 300ng/mL) | 1.4159 (0.9102 - 2.203) | 0.12295 |
|  | **Multivariate (8-gene signature)** |  |
| 8-gene signature (high vs. low risk) | 1.715 (1.054 - 2.791) | **0.0198** |
| Tumour size (> 5cm vs. < 5cm) | 1.1834 (0.4822 - 1.481) | 0.556 |
| Cirrhosis | 2.820 (0.6811 - 11.673) | 0.1527 |
| TNM staging (II & III vs. I) | 1.540 (1.178 - 2.383) | **0.0422** |
| BCLC staging (B & C vs. A & 0) | 1.675 (1.178 - 2.383) | **0.00408** |
| AFP (> 300ng/mL vs. < 300ng/mL) | 1.164 (0.7304 - 1.855) | 0.52297 |
| **TCGA-LIHC** |  |  |
|  | **Univariate** |  |
| 45-gene signature (high vs. low risk) | 1.839 (1.164 - 2.904) | **0.008** |
| 8-gene signature (high vs. low risk) | 2.423 (1.51 - 3.89) | **0.00025** |
| TNM staging (II & III vs. I) | 2.000 (1.533 - 2.608) | **3.14E-07** |
| *TP53* mutation (Yes vs. No) | 2.484 (1.485 - 4.155) | **5.30E-04** |
| *CTNNB1* mutation (Yes vs. No) | 1.894 (1.084 - 3.311) | **0.025** |
| **Multivariate (45-gene signature)** |  |  |
| 45-gene signature (high vs. low risk) | 1.606 (1.014 - 2.542) | **0.03329** |
| TNM staging (II & III vs. I) | 1.879 (1.434 - 2.461) | **4.65E-06** |
| *TP53* mutation (Yes vs. No) | 2.150 (1.271 - 3.635) | **0.0043** |
| *CTNNB1* mutation (Yes vs. No) | 1.553 (0.884 - 2.726) | 0.13 |
| **Multivariate (8-gene signature)** |  |  |
| 8-gene signature (high vs. low risk) | 2.229 (1.388 - 3.580) | **0.000914** |
| TNM staging (II & III vs. I) | 1.865 (1.428 - 2.434) | **4.63E-06** |
| *TP53* mutation (Yes vs. No) | 1.939 (1.158 - 3.246) | **0.012** |
| *CTNNB1* mutation (Yes vs. No) | 1.917 (1.085 - 3.385) | **0.025** |
| **LIRI-JP** |  |  |
| **Univariate** |  |  |
| 45-gene signature (high vs. low risk) | 2.671 (1.361 - 5.240) | **0.00328** |
| 8-gene signature (high vs. low risk) | 3.45 (1.69 - 7.04) | **0.00067** |
| Fibrosis (yes vs. no) | 1.024 (0.8155 - 1.286) | 0.838 |
| Tumour size (> 3cm vs. < 3cm) | 2.415 (1.263 - 4.615) | **0.00764** |
| Portal vein invasion (yes vs. no) | 2.281 (1.712 - 3.039) | **2.00E-08** |
| Hepatic vein invasion (yes vs. no) | 1.822 (1.112 - 2.985) | **0.0173** |
| Bile duct invasion (yes vs. no) | 1.392 (0.987 - 1.963) | 0.059 |
| TNM staging (II & III vs. I) | 2.267 (1.556 - 3.304) | **2.03E-05** |
| **Multivariate (45-gene signature)** |  |  |
| 45-gene signature (high vs. low risk) | 2.201 (1.112 - 4.357) | **0.0236** |
| Tumour size (> 3cm vs. < 3cm) | 1.471 (0.746 - 2.898) | 0.265 |
| TNM staging (II & III vs. I) | 2.009 (1.348 - 2.995) | **0.000617** |
| **Multivariate (45-gene signature)** |  |  |
| 45-gene signature (high vs. low risk) | 2.286 (1.139 - 4.587) | **0.02** |
| Portal vein invasion (yes vs. no) | 2.314 (1.635 - 3.275) | **2.19E-06** |
| Hepatic vein invasion (yes vs. no) | 0.843 (0.462 - 1.539) | 0.579 |
| **Multivariate (8-gene signature)** |  |  |
| 8-gene signature (high vs. low risk) | 2.578 (1.228 - 5.412) | **0.0123** |
| Tumour size (> 3cm vs. < 3cm) | 1.386 (0.698 - 2.752) | 0.3513 |
| TNM staging (II & III vs. I) | 1.878 (1.268 - 2.781) | **0.00167** |
| **Multivariate (8-gene signature)** |  |  |
| 8-gene signature (high vs. low risk) | 2.419 (1.141 - 5.129) | **0.0213** |
| Portal vein invasion (yes vs. no) | 2.057 (1.449 - 2.920) | **5.40E-05** |
| Hepatic vein invasion (yes vs. no) | 0.914 (0.508 - 1.646) | 0.765 |

We next examined the relation of HCC45 to other clinical parameters and observed that it remained highly prognostic after adjustment for tumor size, cirrhosis, fibrosis, TNM stages, Barcelona Clinic Liver Cancer stages, vascular invasion, *TP53/CTNNB1* mutation status and/or alpha-fetoprotein levels in a multivariate Cox regression analysis: GSE14520 hazard ratio [HR] (HR=1.89, P=0.0052), TCGA-LIHC (HR=1.61, P=0.033) and LIRI-JP (HR=2.20, P=0.024) (Table 1). Both cirrhosis and HCC45 harbor complementary prognostic information contributing to elevated HR to 4.59 (95% CI 1.11-19.03, P=0.036) (Table 1).

Gene ontology analysis of HCC45 revealed enrichment of biological pathways linked to hypoxia, metabolism, inflammation and cancer (Fig.S3). Moreover, HCC45 was enriched for targets of several well-known cancer-related transcription factors such as *MYC* and *RUNX1* (Fig.S3) and were functionally connected as a group (protein-protein interaction network enrichment: P<1e-16) (Fig.S4).

### The minimal prognostic 8-gene signature is sufficient and effective in predicting overall survival in HCC

Having identified that HCC45 predicts outcome in HCC, we evaluated the performance of each gene as a prognostic factor (Fig.1). Of the 45 genes, univariate Cox regression analysis revealed that 14, 19 and 28 genes were significantly associated with unfavorable prognosis in GSE14520, TCGA-LIHC and LIRI-JP respectively (Fig.1; Fig.3A). Eight genes (HCC8) were found to confer prognostic information in all HCC cohorts: *CA9, CCL20, CORO1C, CTSC, LDHA, NDRG1, PTP4A3* and *TUBA1B* (Fig.3A; Table S4). Expression of HCC8 not only distinguished tumor from non-tumor sample (Fig.S5) but also increased according to tumor progression (Fig.S6). As expected, the signature successfully identified high-risk patients in full HCC cohorts, GSE14520 (P=0.0001), TCGA-LIHC (P=0.00016) and LIRI-JP (P=0.0003) cohorts, and in patients stratified by tumor stage (Fig.3B). The 5-year disease-free survival rates were also lower in high-risk patients as determined by the 8-gene signature (Fig.S7). Furthermore, as measured by Wald chi-square statistics and log-rank tests, HCC8 is superior in predicting death than HCC45 (Fig.2A; Fig.3B; Table 1).

**Figure 3.**
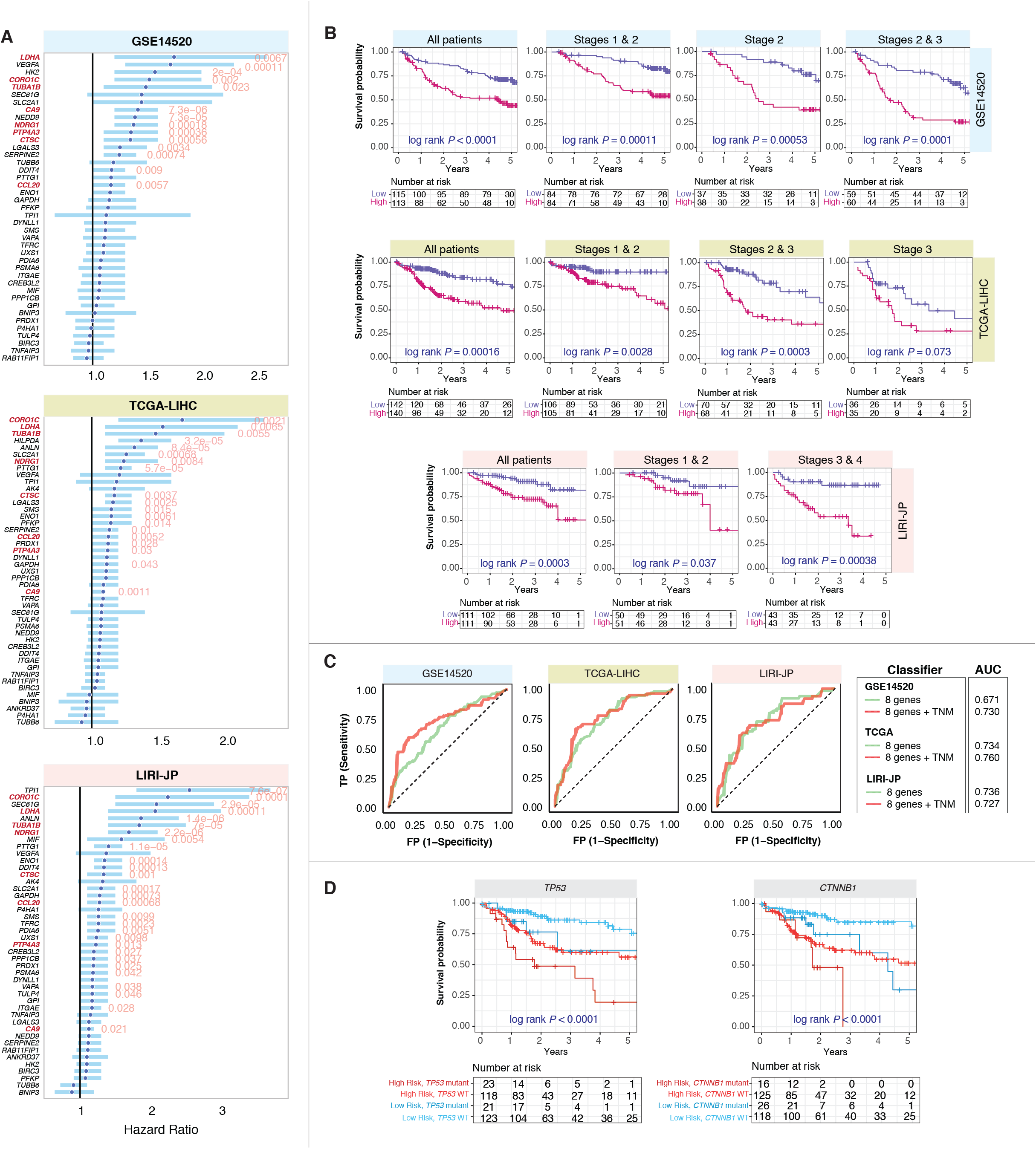
Minimal prognostic 8-gene signature in HCC. **(A)** Forest plots depict Cox proportional hazards analysis on 45 signature genes in three HCC cohorts. Hazard ratios (HR) were denoted as dark blue circles and light blue bars represent 95% confidence interval. Eight genes are consistently prognostic across all three cohorts, thereby constituting the minimal prognostic signature. Significant Wald test P values were indicated in red. Signature genes were highlighted in red. **(B)** The 8-gene signature successfully identified high-risk patients in different TNM stages. Kaplan-Meier plots of overall survival in HCC patients across three cohorts stratified by 8-gene signature into low and high-risk groups. Patients were stratified by the signature as a full cohort, or sub-grouped according to TNM stages. P-values were calculated from the log-rank test. Plots show independence of signature over current staging systems. **(C)** Analysis of specificity and sensitivity of the signature in HCC cohorts with ROC. Plots depict comparison of ROC curves of signature and clinical tumor staging parameters. AUC: area under the curve. TNM: tumor, node, metastasis staging. **(D)** Kaplan-Meier plots depicting combined relation of *TP53* or *CTNNB1* mutation status with the 8-gene signature on overall survival in HCC.

Multivariate models of HCC8 revealed that this reduced gene set sufficiently served as an independent prognostic risk factor: GSE14520 (HR=1.72, P=0.02), TCGA-LIHC (HR=2.23, P=0.0009) and LIRI-JP (HR=2.58, P=0.012) (Table 1). Predictive value for 5-year OS increased when the HCC8 signature was used in combination with TNM staging: GSE14520 AUC=0.67 versus 0.73 and TCGA-LIHC AUC=0.73 versus 0.76 (Fig.3C). When *TP53* or *CTNNB1* mutation status and HCC8 were jointly used, patients with high HCC8 levels and mutant *TP53* (P<0.0001) or *CTNNB1* (P<0.0001) had the lowest chances of survival (Fig.3D).

We calculated risk scores for each patient by taking the sum of Cox regression coefficient for each of the 8 genes multiplied with its corresponding expression value[20]. Risk scores significantly correlated with hypoxia scores in all 3 cohorts, indicating that patients with more hypoxic tumors have higher risk for death as predicted by HCC8 (Fig.S8), overall suggesting that it was sufficient and effective in predicting death.

### Prognosis of the 8-gene signature in 5 other cancers

Hypoxic tumors recruit CD4^+^ Tregs to suppress effector T cell function and promote tumor tolerance[10]. We hypothesized that HCC8 could predict outcome in other solid malignancies in addition to HCC. HCC8 could indeed discriminate between tumor and non-tumor samples in five other cancer types (Fig.S9). Remarkably, our prognostic 8-gene signature derived from HCC was a significant adverse prognostic factor for OS in these five cancer types: head and neck (HR=1.64, P=0.004), renal papillary cell (HR=2.31, P=0.047), lung (HR=1.45, P=0.03), pancreas (HR=1.96, P=0.006) and endometrium (HR=2.33, P=0.003) (Fig.4A; Table S5). Multivariate analysis adjusting for tumor stage revealed that the HCC8 signature retained independent prognostic relevance in these cancers: head and neck (HR=1.66, P=0.003), lung (HR=1.45, P=0.029), pancreas (HR=1.91, P=0.009) and endometrium (HR=1.99, P=0.016) (Table S5). Importantly, while TNM staging was not a reliable predictor of OS in pancreatic cancer on its own, our 8-gene signature successfully predicted high-risk patients when accounting for tumor stage (P=0.009) (Table S5).

**Figure 4.**
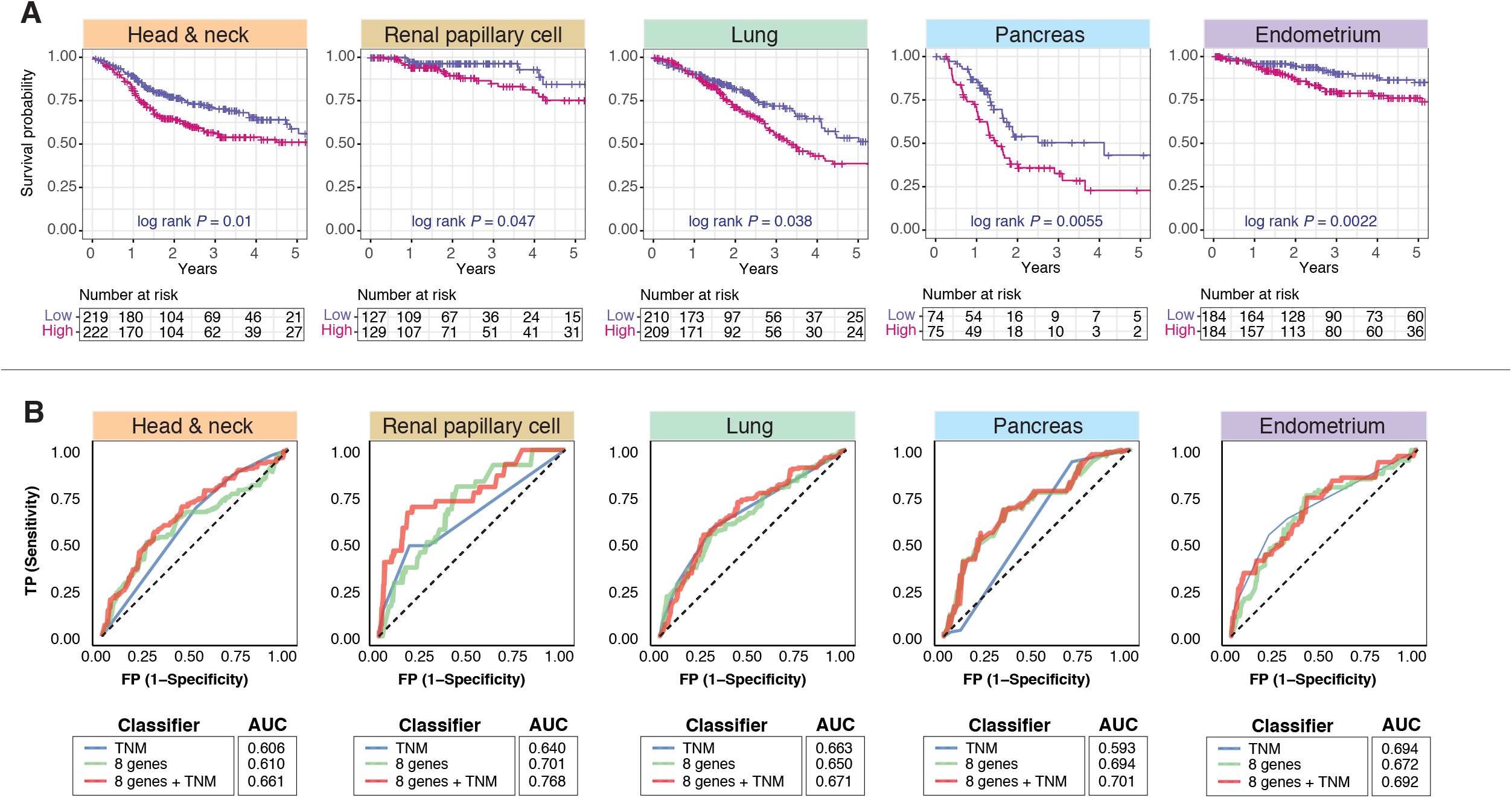
Prognosis of the 8-gene signature in 5 other non-HCC cancers. **(A)** Kaplan-Meier plots of overall survival in patients across multiple cancers stratified into low and high-risk groups using the prognostic 8-gene signature. P-values were obtained from the log-rank test. **(B)** Analysis of specificity and sensitivity of the 8-gene signature in multiple cancers. Plots depict comparison of ROC curves of the 8-gene signature and clinical tumor staging parameters. AUC: area under the curve. TNM: tumor, node, metastasis staging.

The predictive value of the 8-gene signature outperformed the TNM staging system: head and neck (AUC=0.610 versus 0.606), renal papillary cell (AUC=0.701 versus 0.640) and pancreas (AUC=0.694 versus 0.593) (Fig.4B). When used in conjunction with staging information, additive effects on the predictive performance of HCC8 were observed: head and neck (AUC=0.661), renal papillary cell (AUC=0.768), lung (AUC=0.671) and pancreas (AUC=0.701) (Fig.4B). Importantly, risk scores derived from HCC8 showed strong positive trends with tumor hypoxia, suggesting that high risk patients had more aggressive tumors: head and neck (ρ=0.729, P<0.0001), renal papillary cell (ρ=0.766, P<0.0001), lung (ρ=0.856, P<0.0001), pancreas (ρ=0.844, P<0.0001) and endometrium (ρ=0.374, P<0.0001) (Fig.S10).

*TP53, KRAS* and *CDKN2A* mutations are commonly observed in lung cancer[21]. *CDKN2A* (P=0.03), but not *TP53* (P=0.2) or *KRAS* (P=0.4), was significantly associated with poor prognosis (Table S5). When used in combination with our HCC8 signature, patients with high HCC8 score and mutant *CDKN2A* had ~30% higher chances for death at 2 years than patients with low HCC8 and wild-type *CDKN2A* (Fig.S11).

## Discussion

We developed a novel gene signature that predicts overall survival in six cancers. This represents a significant step forward in the prognostic pathway since pan-cancer genetic signatures are limited at best. Capitalizing two biological phenomena of solid tumors, hypoxia and immune cells infiltration, our signature can be applied across multiple cancers, suggesting that a high degree of commonality exists in host immune response within a hypoxic tumor microenvironment. Signature genes may provide insights into biological features of individual tumors as they are implicated in oncogenic processes, i.e. pH regulation, invasion, cell proliferation, adhesion and migration, glycolysis and inflammation[22–26]. Combining the RNA gene signature with DNA somatic mutations simultaneously improved its prognostic capabilities, indicating that signature analysis at two macromolecular levels offers the opportunity for multiple levels of drug targeting. High-risk patients, as predicted by our signature, that concurrently have *TP53* mutations suggests a role for hypoxia in promoting genomic instability and aberrant DNA damage repair.

Canonical tumor staging parameters are useful, but they do not offer additional resolution for discriminating patients with similarly-staged malignant grades. Especially for patients with stage 1 cancer, the signature allows the incorporation of hypoxia information to identify patients that are more at risk of progressing to advanced stages and developing lethal metastasis. Additionally, we were able to differentiate tumor from non-tumor samples using the signature, suggesting that it offers additional information on how transformed cells differ from normal cells. It is interesting to speculate that the signature could be used in diagnostics since non-transformed cells would have a distinct expression profile and a deviation from this profile could indicate the onset of oncogenesis.

Tumor hypoxia has wide-ranging effects causing metabolic alteration, angiogenesis, metastasis and immune suppression[27]. Significant crosstalk exists between hypoxia and antitumor immune functions where tumor hypoxia contributes to attenuated antitumor responses[10]. We observed a significant positive correlation between Treg and hypoxia gene expressions, supporting the notion that immunosuppression is higher in patients with more hypoxic tumors (Fig.S1C,D). Cancerous cells are protected through immunosuppressive functions enhanced by tumor hypoxia and together, they contribute to chemotherapy resistance[28]. Hence, to improve patient response to treatment, there is an urgent need to incorporate hypoxic tumor assessments in clinic for an effective patient-centric strategy. Through personalized therapy, patients with more hypoxic tumors would benefit from neoadjuvant treatment using hypoxia-modifying drugs to improve response to immunotherapy and chemotherapy[14].

Elevation of Tregs and exhausted CD8^+^ T cells in tumors offers another opportunity for therapeutic intervention. Exhausted CD8^+^ T cells are overrepresented among tumor-infiltrating CD8^+^ T cells[8]. These cells have reduced levels of cytotoxic markers and high expression of the *PDCD1* exhaustion marker[8]. Hence, patients with high signature scores may likely have increased Treg and exhausted CD8^+^ T cell densities. These patients could benefit from therapy with the PD1 antibody as anti-PD1 treatment can rejuvenate exhausted CD8^+^ T cells[29]. When used in combination with first-line treatments, this may dramatically improve patient prognosis.

Although prospective validation is warranted, we consider our results as supporting the implications of tumor’s hypoxic and immunologic microenvironment in influencing patients’ prognosis and potentially their response to treatment. Multi-cohort validations incorporating large sample size (n=2,712) confirmed that the 8-gene signature is robust, clinically actionable and has potential to radically change how we determine prognosis and guide therapy.

## Methods

Detailed methods are available in supplementary information.

## Supplementary methods

### Study cohorts

HCC data sets (n=839) used in this study were GSE14520 (n=242) from the Liver Cancer Institute, LIRI-JP (n=226) from the International Cancer Genome Consortium (ICGC) database and TCGA-LIHC (n=371) from the Cancer Genome Atlas (TCGA) Research Network[13]. Full clinical characteristics of HCC patients were listed in Table S2. Among these patients, underlying etiology differed between the cohorts with GSE14520 comprising of patients with HBV, while LIRI-JP and TCGA-LIHC consisting of patients with HBV, HCV and/or non-alcoholic steatohepatitis.

Gene expression data for 24 other cancer types (n=20,241) generated by TCGA Research Network[13] were downloaded from Broad Institute GDAC Firehose. Illumina HiSeq rnaseqv2 Level 3 RSEM normalized gene expression profiles were converted to log_2_(x + 1) scale. Expression values of genes associated with tumor-infiltrating T cells were corrected for levels of infiltration by dividing expression values of each gene with *CD3D* values.

### Differential expression analysis

To determine differentially expressed genes between tumor and adjacent normal liver tissues in the GSE14520 cohort, the Bayes method and linear model were employed using the R package limma. P-values were adjusted using the false discovery rate controlling procedure of Benjamini-Hochberg. Genes with log_2_ fold change of > 1 and adjusted P-values < 0.05 were considered significant.

### Gene signature and risk scores

Expression scores for hypoxia, tumor-infiltrating T cells, 45-gene signature and 8-gene signature were calculated for each patient in all the cohorts by taking the average log_2_ expression values. Nonparametric Mann-Whitney-Wilcoxon test was used to compare the distribution of hypoxia scores in tumor and adjacent non-tumor samples. Risk scores for each patient were determined by taking the sum of Cox regression coefficient for each of the signature genes multiplied with its corresponding expression value. Nonparametric Spearman’s rank correlation was employed to assess the relationship of gene scores and risk scores with tumor hypoxia (hypoxia score).

### Survival analyses

We employed the Cox proportional hazards model to investigate the association between patient survival duration and one or more risk factors, e.g. 45-gene or 8-gene score, tumor stage and other clinical variables. Univariate analysis was performed to determine the influence of individual risk factors on overall survival. As multiple covariates can potentially influence patient prognosis, multivariate analyses were performed by including risk factors that were significantly associated with overall survival identified in univariate analysis (P < 0.05). Hazard ratios (HR) were determined from Cox models with HR greater than one indicating that a covariate was positively associated with event probability (increased hazard) and negatively associated with survival length. Cox regression analyses were performed using the R survival and survminer packages. Proportional hazards assumption was supported by a non-significant relationship between scaled Schoenfeld residuals and time using the R survival package. In addition, Kaplan-Meier and log-rank tests were used in univariate analyses of the gene signatures in relation to patient survival and were performed using the survival and survminer packages. Patients were median-dichotomized into low and high-risk groups based on mean expression scores of signature genes and Kaplan-Meier estimates of survival probability over time were generated. Difference between high and low-risk groups were tested using the log-rank test.

Time-dependent receiver operating characteristic (ROC) curve analysis was used to assess the predictive performance of both 45-gene and 8-gene signatures in comparison with standard tumor staging parameters. The R survcomp package was employed to compute time-dependent ROC curves. ROC curves depicted true positive rates (sensitivity) versus false positive rates (1-specificity). The area under the curve (AUC) is a measure of how well the gene signatures can predict patient survival where AUC ranges from 0.5 (for an uninformative marker) to 1 (for a perfect predictive marker).

### Visualization of sample distance in the reduced 2-dimensional space

To determine whether the gene signatures (45-gene or 8-gene signature) can distinguish tumor and non-tumor samples, multidimensional scaling (MDS) analyses were performed using the R vegan package to visualize samples’ distance in the reduced 2-dimensional space. Euclidean genetic distances between each sample were investigated by MDS ordination. Permutational multivariate analysis of variance (PERMANOVA) was employed to test for differences between tumor and non-tumor samples.

### Biological enrichment and protein interaction network analyses

Analysis of biological pathway enrichment on the 45-gene set was conducted using GeneCodis against the Kyoto Encyclopedia of Genes and Genomes (KEGG) and Gene Ontology (GO) databases. The Enrichr tool was used to identify transcription factors from the ChEA database that are potential regulators of the 45 genes. Functional protein association network of the 45 genes was determined using the STRING database.

### Somatic mutation identification

Level 3 mutation datasets were downloaded from GDAC. Kaplan-Meier analysis and log-rank tests were employed to determine the association of somatic mutations in combination with the 45-gene or 8-gene signatures, on overall survival.

All graphs were generated using the ggplot2 package in R. Heatmap was generated using the R pheatmap package.

## Acknowledgements

A special thank you to Graham Foster for his critical insights and help on manuscript preparation.

## Authors contribution

WHC and AGL designed the study and analyzed the data. All authors interpreted the data. WHC and AGL wrote the initial manuscript draft. All authors revised the manuscript and approved the final version.

---

**Figure S1.**
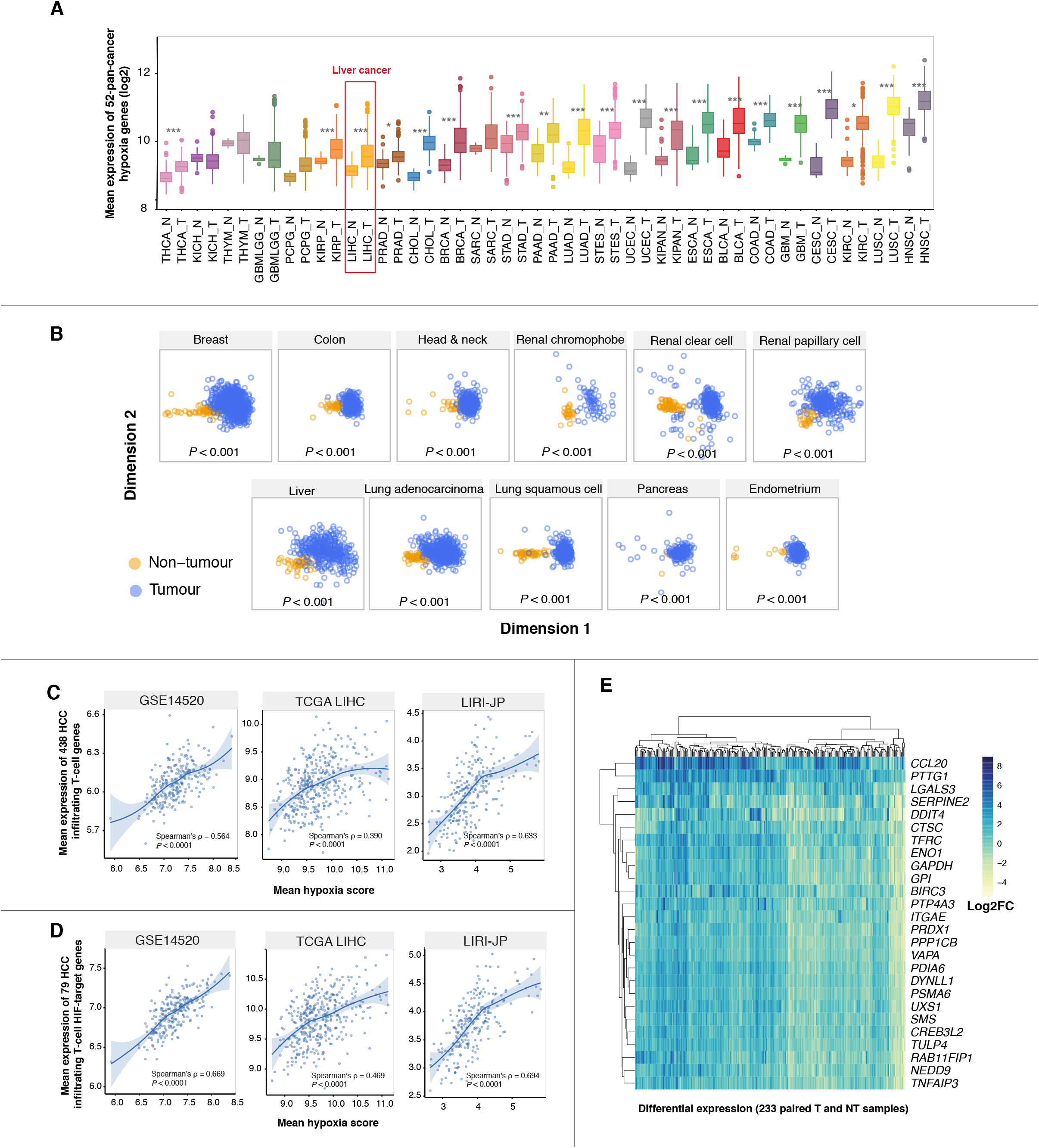
Genes associated with tumor hypoxia and T-cell infiltration. (A) Box plot depicts hypoxia score between tumor (T) and non-tumor(N) samples across 25 cancer datasets obtained from TCGA. Hypoxia scores were estimated by obtaining the mean expression (log2) of 52 hypoxia genes reported by Buffa et al. (2010). Cancer types were sorted on the basis of smallest to largest median hypoxia score in tumor samples. Distribution of hypoxia scores for T and NT samples for each cancer was compared using the Mann-Whitney-Wilcoxon test. Asterisks represent significant P values: 
<0.01, **<0.001 and ***<0.0001. TCGA abbreviations were used to represent cancer types; refer to Table S1. (B) Ordination plots of multidimensional scaling analysis of the 52 hypoxia signature genes using Euclidean distances revealed significant separation of tumor (T) and non-tumor (NT) samples represented in a 2-dimensional space. Axes represent the first and second dimension. The distinction of T and NT was confirmed by permutational multivariate analysis of variance (PERMANOVA) tests. Analysis was performed using the metaMDS and adonis function of the R vegan package. (C and D) Significant positive correlation between HCC-infiltrating gene expression and tumor hypoxia. Expression of both (C) 438 gene set (HCC infiltrating T-cells) and (D) 79 gene set (HIF-target genes associated with HCC infiltrating T-cells) positively correlated with tumor hypoxia as determined from the Buffa hypoxia gene signature. (E) Heatmap depicts differential expression value of 26 HCC-upregulated genes in 233 tumor and non-tumor paired samples from the training cohort. Of the 79 HIF-target genes associated with HCC infiltrating T-cells, 26 are at least 1.5-fold significantly upregualted.

**Figure S2.**
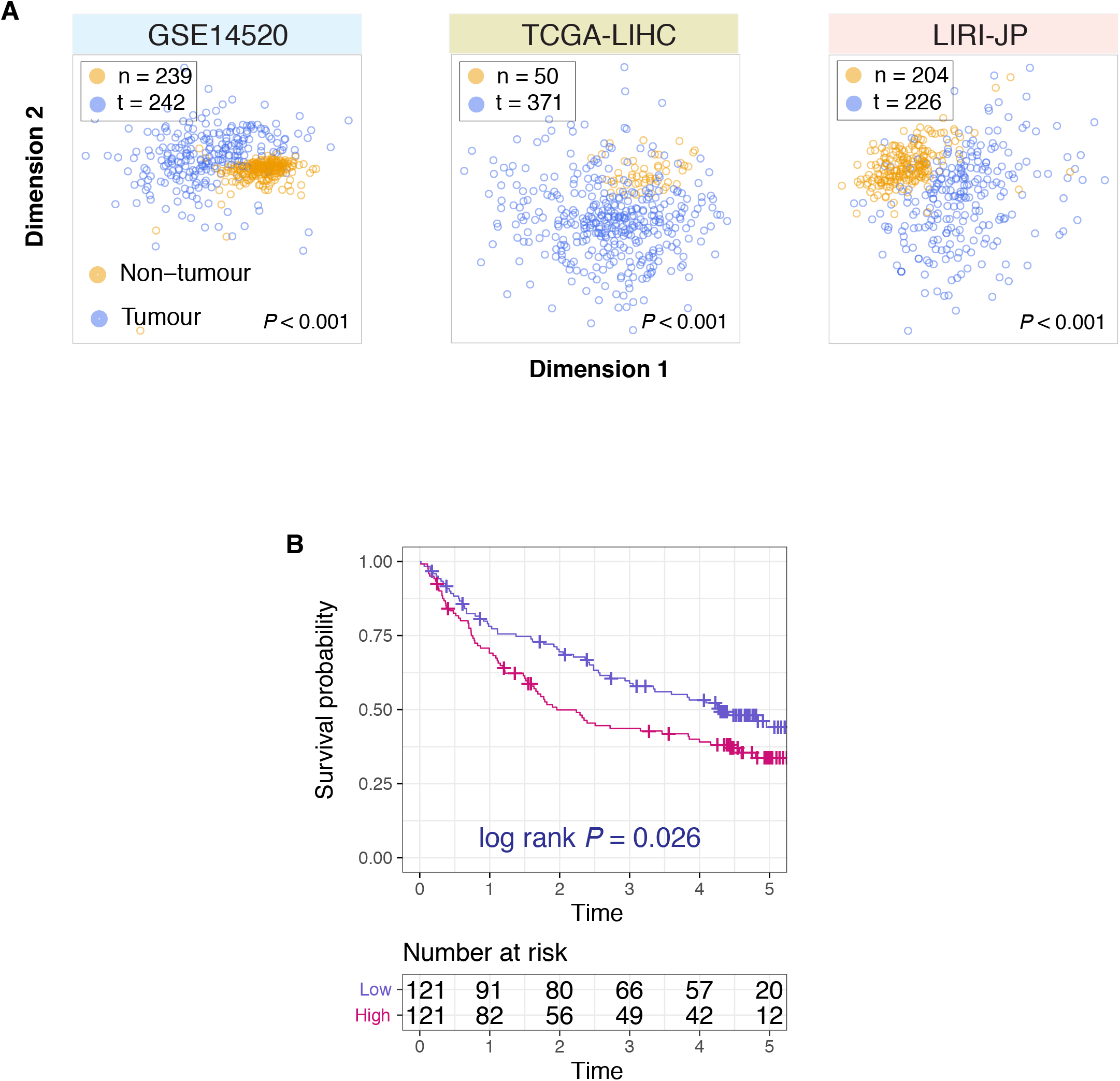
Multidimensional scaling analysis of the 45-gene signature and disease-free survival analysis. **(A)** Ordination plots of multidimensional scaling analysis the signature in HCC cohorts using Euclidean distances revealed significant separation of tumor (T) and non-tumor (NT) samples represented in a 2-dimensional space. Axes represent the first and second dimension. The distinction of T and NT was confirmed by PERMANOVA tests. **(B)** Kaplan-Meier plot of disease-free survival in HCC patients from the GSE14520 cohort stratified into low- and high-risk groups using the 45-gene signature. Disease-free survival is defined as the time from surgery to recurrence, death from any cause or distant metastasis. P-values are calculated from the log-rank test.

**Figure S3.**
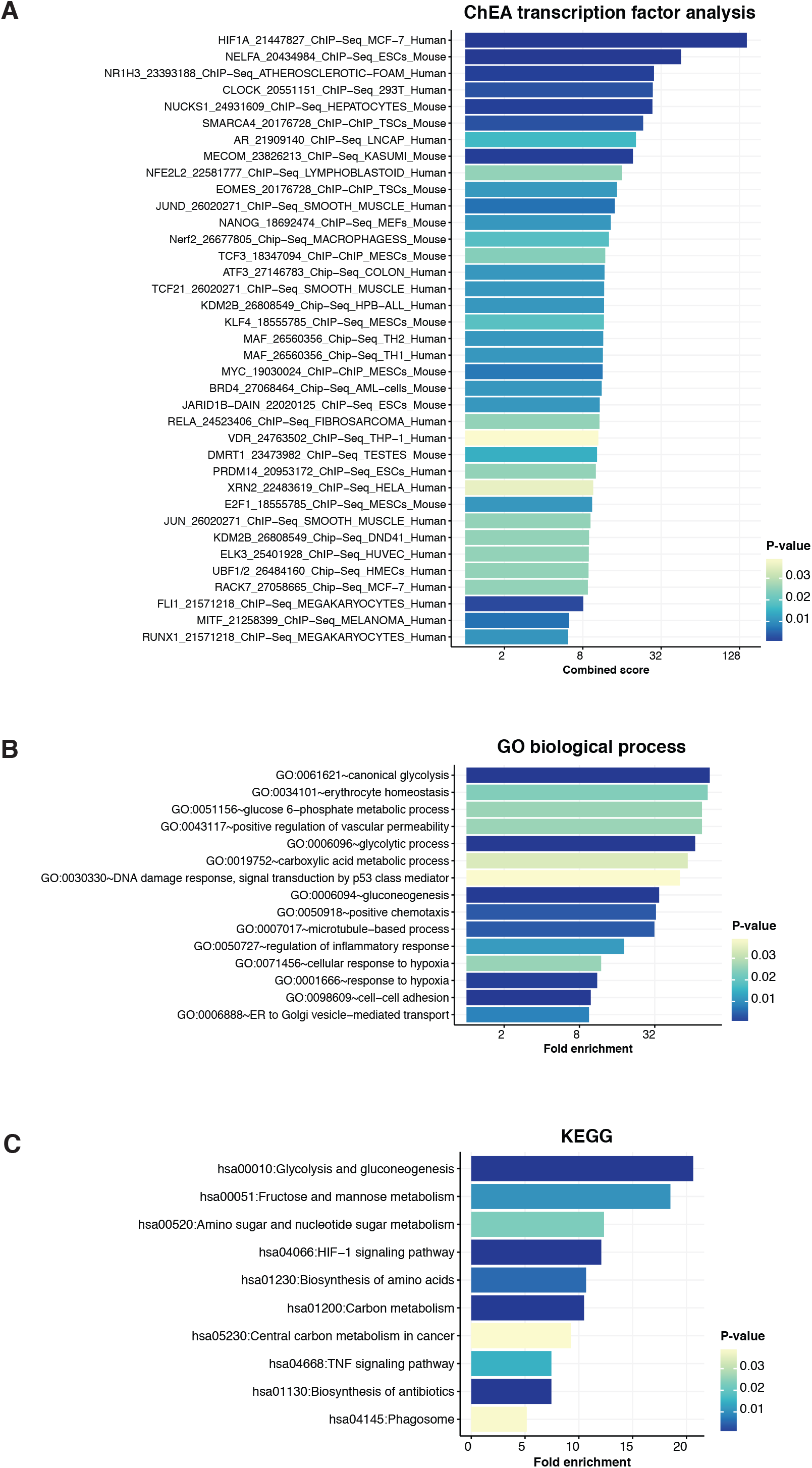
Biological functions associated with the 45-gene signature revealed enrichments of pathways associated with hypoxia, metabolism and cancer. **(A)** Transcription factors and histone modifiers that are potential regulators of the 45 genes. **(B)** Enrichment of GO biological processes. **(C)** Enrichment of KEGG ontologies.

**Figure S4.**
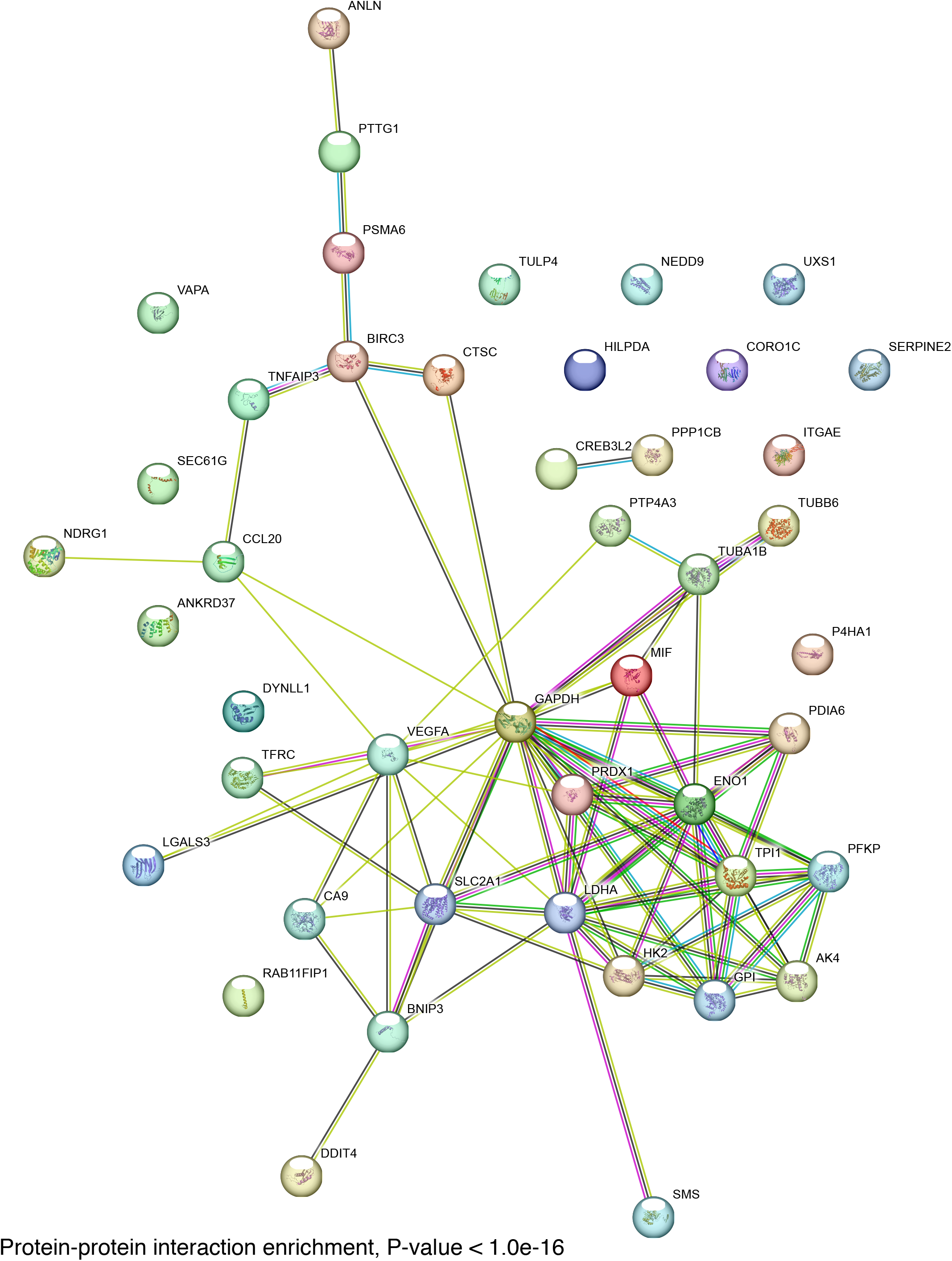
Protein-protein interaction (PPI) networks associated with the 45-gene signature. As determined by STRING (version 10.5), PPI enrichment was significant (P < 1e-16) indicating that the proteins are biologically connected as a group.

**Figure S5.**
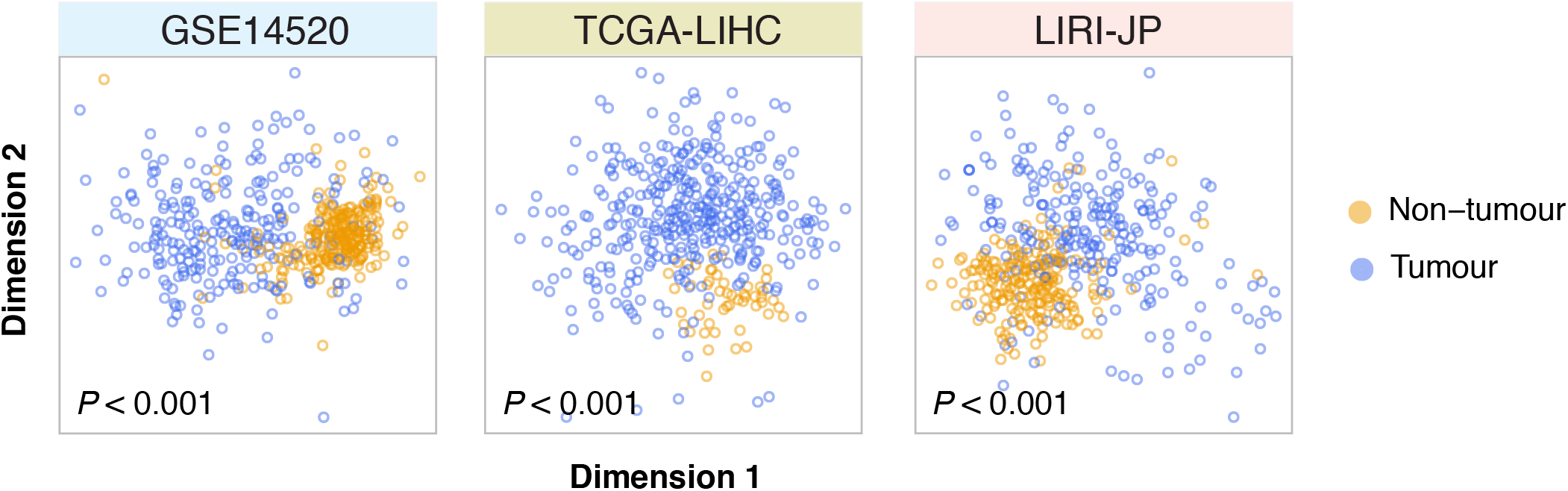
Ordination plots of multidimensional scaling analysis of the 8-gene signature in HCC cohorts using Euclidean distances revealed significant separation of tumor (T) and non-tumor (NT) samples represented in a 2-dimensional space. Axes represent the first and second dimension. The distinction of T and NT was confirmed by PERMANOVA tests.

**Figure S6.**
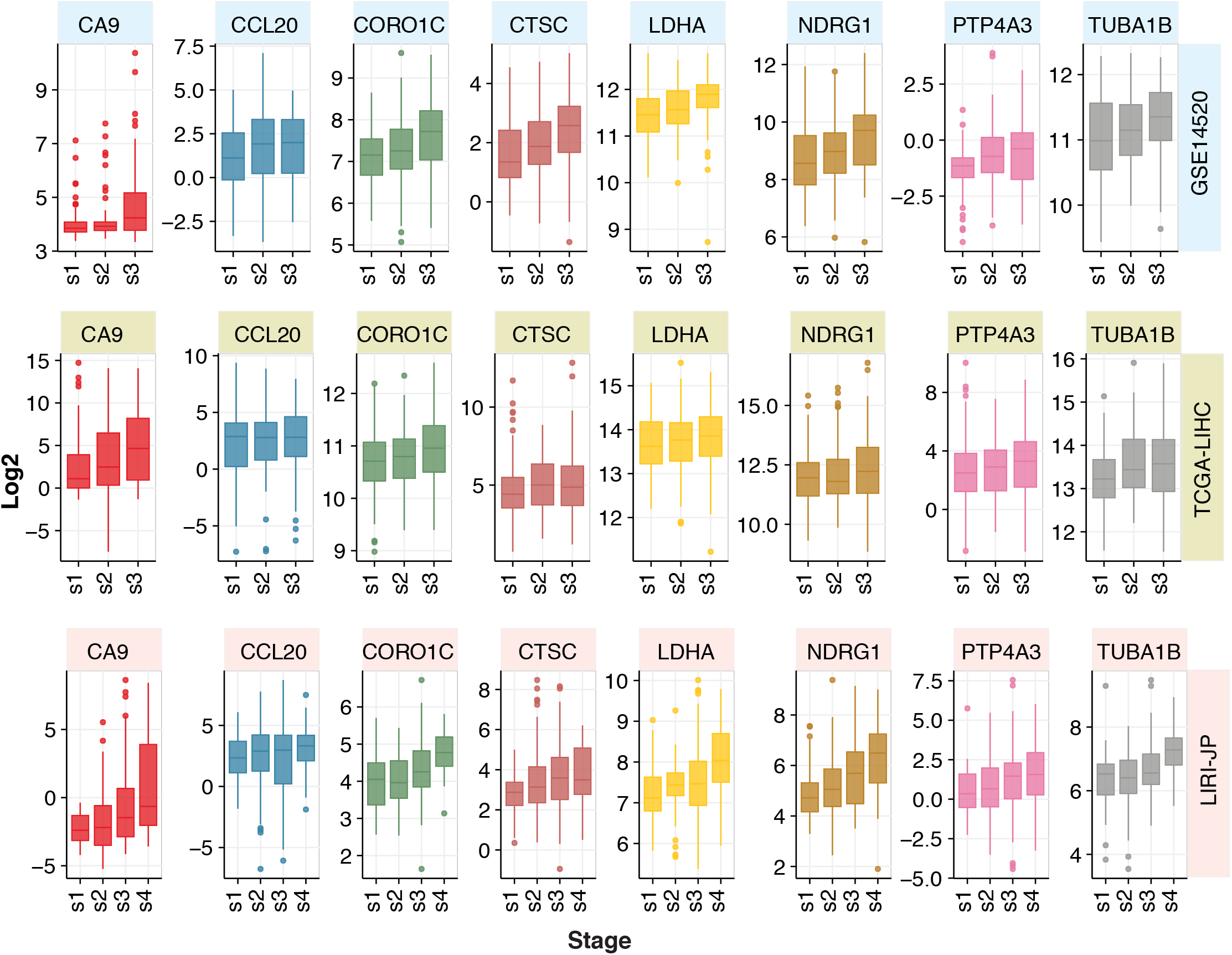
Expression distribution of genes from the 8-gene signature according to tumor staging in three HCC cohorts. Expression of genes increased with tumor staging in HCC patients.

**Figure S7.**
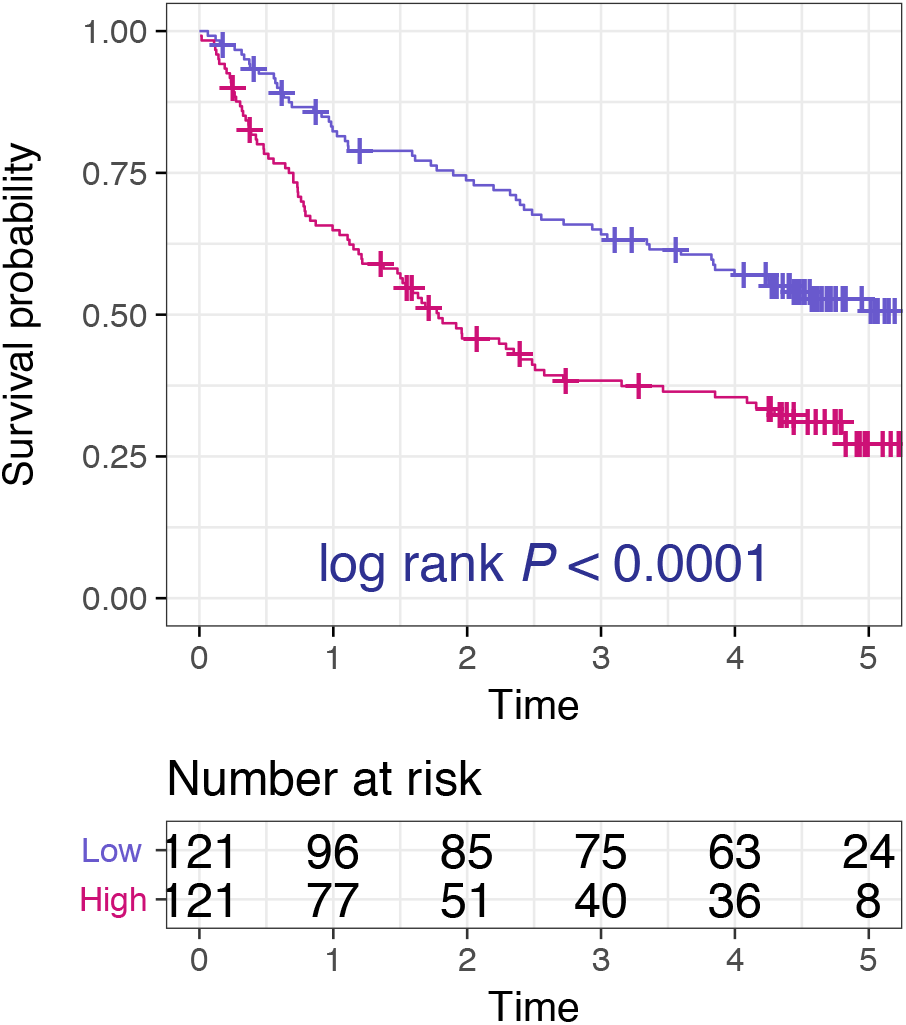
Kaplan-Meier plot of disease-free survival in HCC patients from the GSE14520 cohort stratified into low- and high-risk groups using the 8-gene signature. Disease-free survival is defined as the time from surgery to recurrence, death from any cause or distant metastasis. P-values are calculated from the log-rank test.

**Figure S8.**
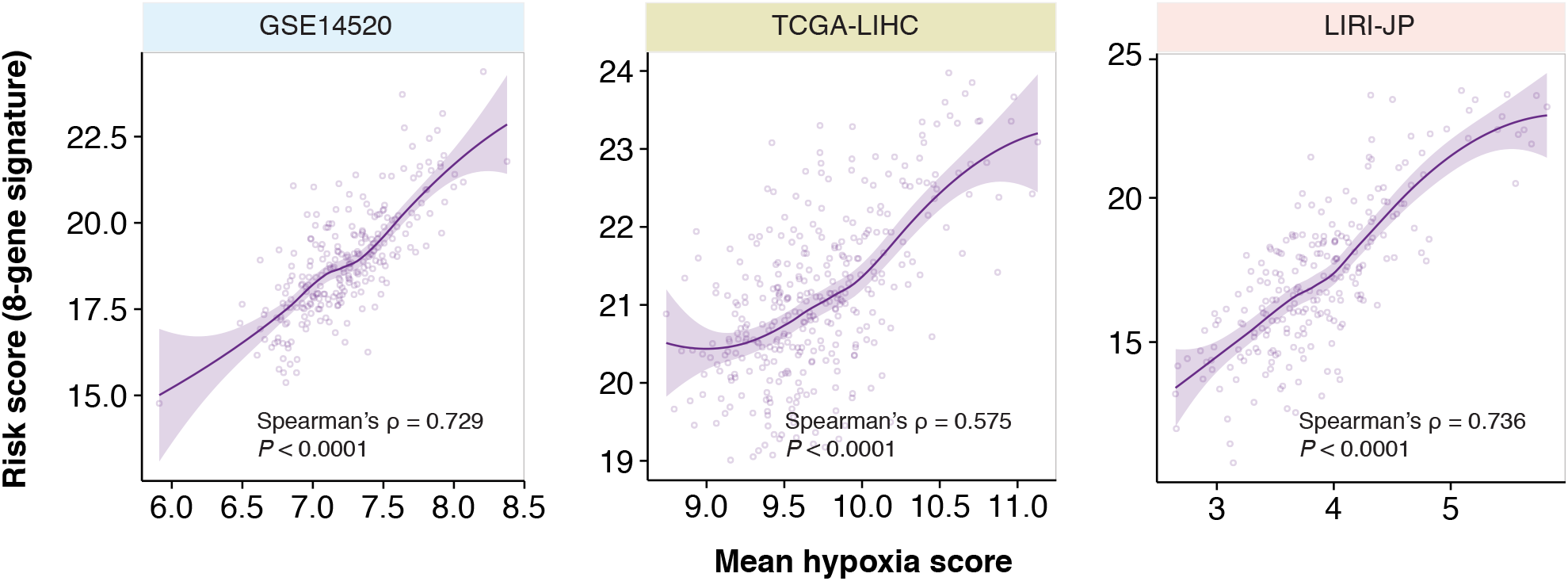
Correlation of risk scores, as determined using the 8-gene signature, and hypoxia scores in HCC patients. Significant positive correlation between patient survival risk scores (refer to methods) derived from the 8-gene signature and tumor hypoxia in HCC cohorts.

**Figure S9.**
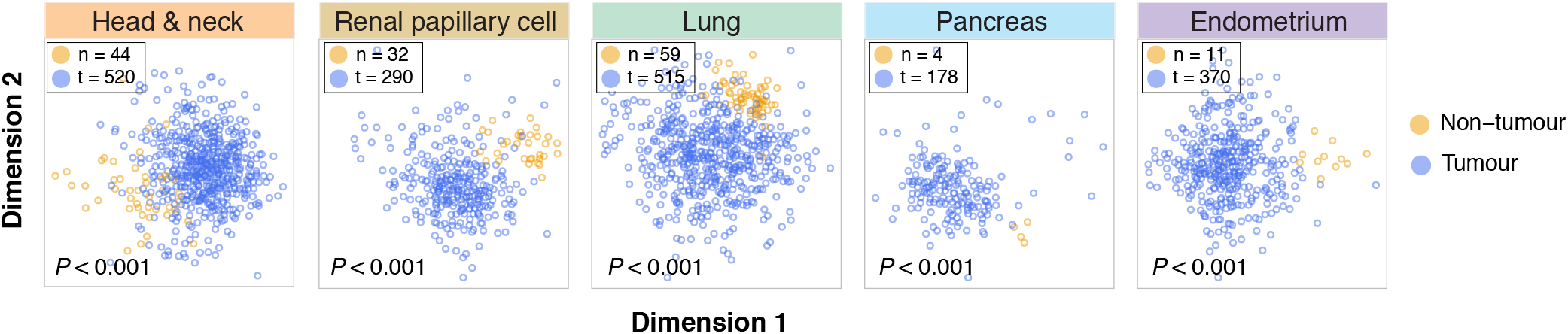
Ordination plots of multidimensional scaling analysis of the 8-gene signature in cancers using Euclidean distances revealed significant separation of tumor (T) and non-tumor (NT) samples represented in a 2-dimensional space. Axes represent the first and second dimension. The distinction of T and NT was confirmed by PERMANOVA tests.

**Figure S10.**
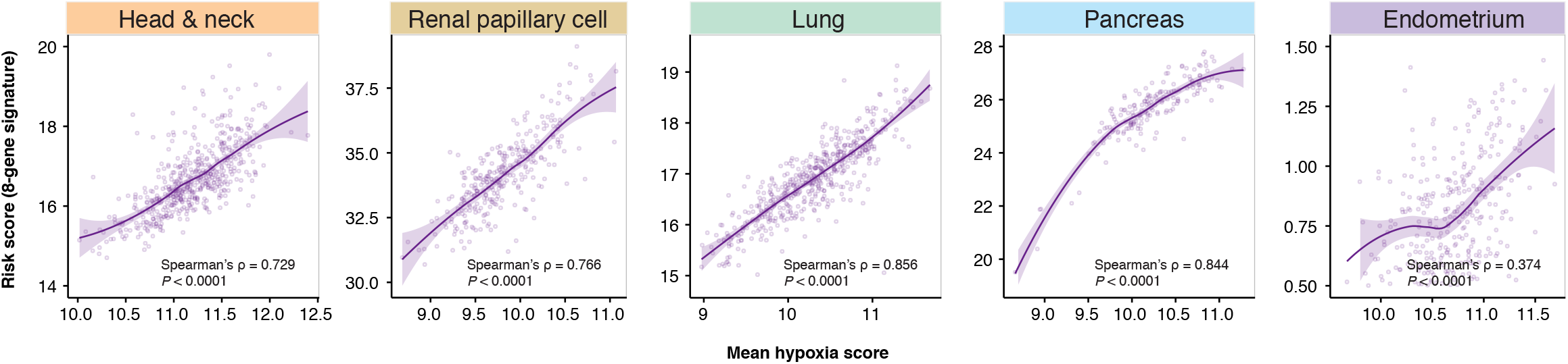
Correlation of risk scores, as determined using the 8-gene signature, and hypoxia scores in other cancers. Significant positive correlation between patient survival risk scores (refer to methods) derived from the 8-gene signature and tumor hypoxia in cancers.

**Figure S11.**
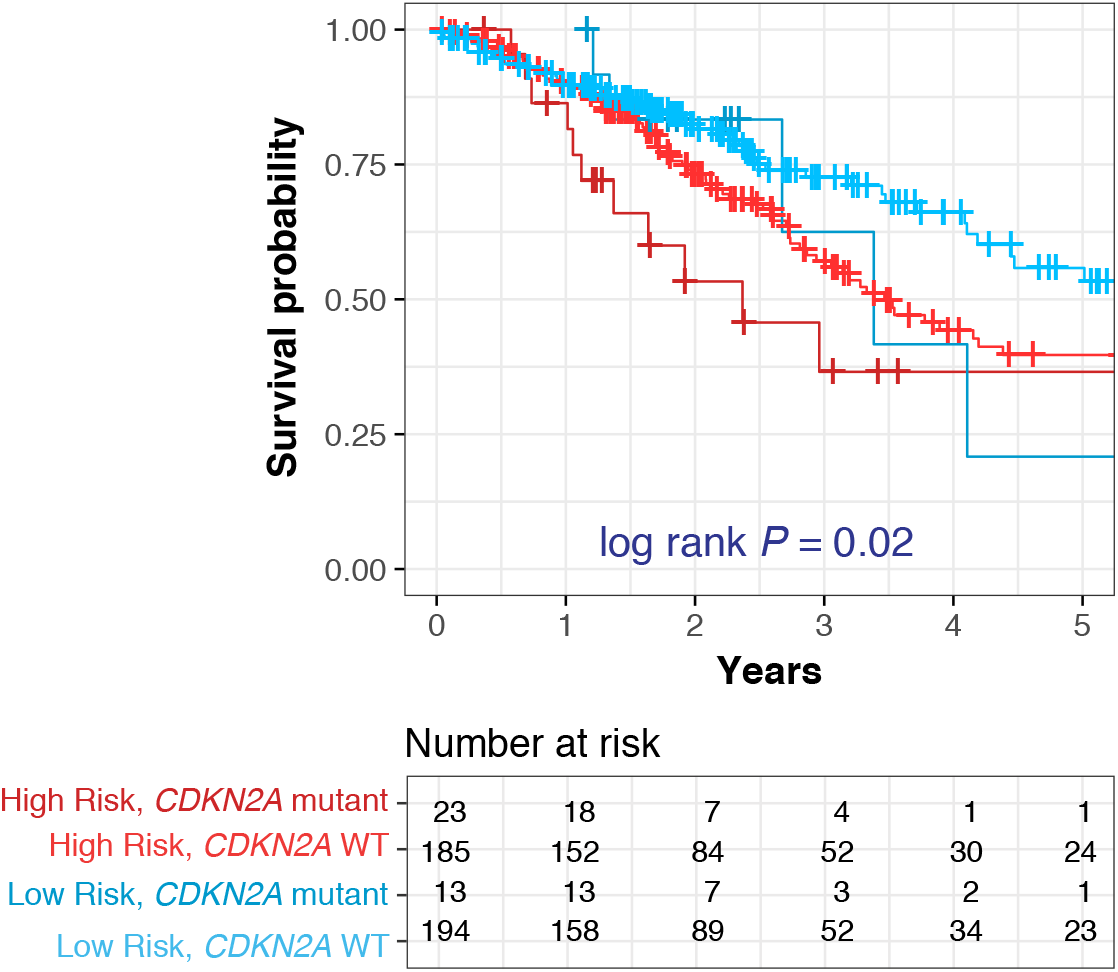
Kaplan-Meier plot depicting combined relation of *CDKN2A* mutation status with the 8-gene signature on overall survival in lung cancer.

**Table S1.** Abbreviations and number of tumour and non-tumour samples in TCGA cancers.

| **Cohort (TCGA abbreviations)** | **Non-tumour #** | **Tumour #** | **Description** |
| --- | --- | --- | --- |
| BLCA | 19 | 408 | Bladder Urothelial Carcinoma |
| BRCA | 112 | 10939 | Breast invasive carcinoma |
| CESC | 3 | 304 | Cervical squamous cell carcinoma and endocervical adenocarcinoma |
| CHOL | 9 | 36 | Cholangiocarcinoma |
| COAD | 41 | 285 | Colon adenocarcinoma |
| ESCA | 11 | 184 | Esophageal carcinoma |
| GBM | 5 | 153 | Glioblastoma multiforme |
| GBMLGG | 5 | 669 | Glioma |
| HNSC | 44 | 520 | Head and Neck squamous cell carcinoma |
| KICH | 25 | 66 | Kidney Chromophobe |
| KIPAN | 129 | 889 | Pan-kidney cohort |
| KIRC | 72 | 533 | Kidney renal clear cell carcinoma |
| KIRP | 32 | 290 | Kidney renal papillary cell carcinoma |
| LIHC | 50 | 371 | Liver hepatocellular carcinoma |
| LUAD | 59 | 515 | Lung adenocarcinoma |
| LUSC | 51 | 501 | Lung squamous cell carcinoma |
| PAAD | 4 | 178 | Pancreatic adenocarcinoma |
| PCPG | 3 | 179 | Pheochromocytoma and Paraganglioma |
| PRAD | 52 | 497 | Prostate adenocarcinoma |
| SARC | 2 | 259 | Sarcoma |
| STAD | 35 | 415 | Stomach adenocarcinoma |
| STES | 46 | 599 | Stomach and Esophageal carcinoma |
| THCA | 59 | 501 | Thyroid carcinoma |
| THYM | 2 | 120 | Thymoma |
| UCEC | 11 | 370 | Uterine Corpus Endometrial Carcinoma |

**Table S2.** Clinical characteritics of HCC patients (n = 839)

| **Characteristic** |  | **GSE14520** | **TCGA-LIHC** | **LIRI-JP** |
| --- | --- | --- | --- | --- |
| Patient number | Total | 242 | 371 | 226 |
|  | Male | 211 (87%) | 252 (68%) | 166 (73%) |
|  | Female | 31 (13%) | 119 (32%) | 60 (27%) |
| Age (years) | Range | 21 - 77 | 16 - 90 | 31 - 89 |
|  | Median | 50 | 61 | 68 |
| Viral etiology | Hepatitis B | 242 (100%) | 143 (39%) | 59 (26%) |
|  | Hepatitis C |  | 44 (12%) | 120 (53%) |
|  | Hepatitis B & C |  | 87 (24%) | 4 (2%) |
|  | Negative |  | 7 (2%) | 43 (19%) |
|  | Not available |  | 90 (23%) |  |
| Edmondson grade | I |  |  | 28 (12%) |
|  | II |  |  | 156 (69%) |
|  | III |  |  | 21 (9%) |
|  | IV |  |  | 1 (0.4%) |
|  | Not available | 242 (100%) | 371 (100%) | 20 (9%) |
| Tumour size | Small (< 5cm) | 153 (63%) |  | 174 (77%) |
|  | Large (> 5cm) | 89 (37%) |  | 52 (23%) |
|  | Not available |  | 371 (100%) |  |
| Fibrosis | Negative | 19 (8%) | 76 (20%) | 21 (9%) |
|  | Portal fibrosis (stages 1 -2) |  | 31 (8%) | 73 (32%) |
|  | Fibrous septa (stages 3 - 4) |  | 30 (8%) | 132 (58%) |
|  | Nodular formation (stage 5) |  | 9 (2%) |  |
|  | Cirrhosis (stage 6) | 223 (92%) | 72 (19%) |  |
|  | Not available |  | 153 (41%) |  |
| Alcohol consumption | Yes |  | 118 (32%) | 86 (38%) |
|  | No |  |  | 128 (57%) |
|  | Not available | 242 (100%) | 253 (68%) | 12 (5%) |
| Smoker | Yes |  | 13 (4%) | 122 (54%) |
|  | No |  |  | 94 (42%) |
|  | Not available | 242 (100%) | 358 (96%) | 10 (4%) |
| AFP | Low (< 300ng/mL) | 128 (53%) | 218 (59%) |  |
|  | High (> 300ng/mL) | 110 (45%) | 66 (18%) |  |
|  | Not available | 4 (2%) | 87 (23%) | 226 (100%) |
| BCLC stage | 0 | 20 (8%) |  |  |
|  | A | 152 (63%) |  |  |
|  | B | 24 (10%) |  |  |
|  | C | 29 (12%) |  |  |
|  | Not available | 17 (7%) | 371 (100%) | 226 (100%) |
| TNM stage | I | 96 (40%) | 144 (39%) | 35 (15%) |
|  | II | 78 (32%) | 67 (18%) | 101 (45%) |
|  | III | 51 (21%) | 71 (19%) | 68 (30%) |
|  | IV |  |  | 18 (8%) |
|  | Not available | 17 (7%) | 89 (24%) | 4 (2%) |

**Table S3.** The 45-gene classifier associated with tumour hypoxia and T-cell infiltration.

| Gene Symbol | Description |
| --- | --- |
| AK4 | adenylate kinase 4 |
| ANKRD37 | ankyrin repeat domain 37 |
| ANLN | anillin actin binding protein |
| BIRC3 | baculoviral IAP repeat containing 3 |
| BNIP3 | BCL2 interacting protein 3 |
| CA9 | carbonic anhydrase 9 |
| CCL20 | C-C motif chemokine ligand 20 |
| CORO1C | coronin 1C |
| CREB3L2 | cAMP responsive element binding protein 3 like 2 |
| CTSC | cathepsin C |
| DDIT4 | DNA damage inducible transcript 4 |
| DYNLL1 | dynein light chain LC8-type 1 |
| ENO1 | enolase 1 |
| GAPDH | glyceraldehyde-3-phosphate dehydrogenase |
| GPI | glucose-6-phosphate isomerase |
| HILPDA | hypoxia inducible lipid droplet associated |
| HK2 | hexokinase 2 |
| ITGAE | integrin subunit alpha E |
| LDHA | lactate dehydrogenase A |
| LGALS3 | galectin 3 |
| MIF | macrophage migration inhibitory factor |
| NDRG1 | N-myc downstream regulated 1 |
| NEDD9 | neural precursor cell expressed, developmentally down-regulated 9 |
| P4HA1 | prolyl 4-hydroxylase subunit alpha 1 |
| PDIA6 | protein disulfide isomerase family A member 6 |
| PFKP | phosphofructokinase, platelet |
| PPP1CB | protein phosphatase 1 catalytic subunit beta |
| PRDX1 | peroxiredoxin 1 |
| PSMA6 | proteasome subunit alpha 6 |
| PTP4A3 | protein tyrosine phosphatase type IVA, member 3 |
| PTTG1 | pituitary tumor-transforming 1 |
| RAB11FIP1 | RAB11 family interacting protein 1 |
| SEC61G | Sec61 translocon gamma subunit |
| SERPINE2 | serpin family E member 2 |
| SLC2A1 | solute carrier family 2 member 1 |
| SMS | spermine synthase |
| TFRC | transferrin receptor |
| TNFAIP3 | TNF alpha induced protein 3 |
| TPI1 | triosephosphate isomerase 1 |
| TUBA1B | tubulin alpha 1b |
| TUBB6 | tubulin beta 6 class V |
| TULP4 | tubby like protein 4 |
| UXS1 | UDP-glucuronate decarboxylase 1 |
| VAPA | VAMP associated protein A |
| VEGFA | vascular endothelial growth factor A |

**Table S4.** The minimal prognostic 8-gene classifier.

| Gene Symbol | Description |
| --- | --- |
| CA9 | carbonic anhydrase 9 |
| CCL20 | C-C motif chemokine ligand 20 |
| CORO1C | coronin 1C |
| CTSC | cathepsin C |
| LDHA | lactate dehydrogenase A |
| NDRG1 | N-myc downstream regulated 1 |
| PTP4A3 | protein tyrosine phosphatase type IVA, member 3 |
| TUBA1B | tubulin alpha 1b |

**Table S5.** Univariate and multivariate Cox proportional hazards analysis of risk factors associated with overall survival in multiple cancers.

|  | Univariate |  | Multivariate |  |
| --- | --- | --- | --- | --- |
|  | Hazard Ratio (95% CI) | P-value | Hazard Ratio (95% CI) | P-value |
| **Head and neck** |  |  |  |  |
| 8-gene signature (high vs. low risk) | 1.64 (1.166 - 2.308) | **4.70E-03** | 1.658 (1.178 - 2.322) | **3.74E-03** |
| TNM staging (II & III vs. I) | 1.466 (1.188 - 1.808) | **3.53E-04** | 1.475 (1.194 - 1.820) | **3.05E-04** |
| *TP63* mutation (Yes vs. No) | 1.046 (0.658 - 1.663) | 0.85 |  |  |
| *CDKN2A* mutation (Yes vs. No) | 1.318 (0.892 - 1.947) | 0.17 |  |  |
| *PIK3CA* mutation (Yes vs. No) | 1.261 (0.785 - 2.026) | 0.34 |  |  |
| **Renal papillary cell** |  |  |  |  |
| 8-gene signature (high vs. low risk) | 2.311 (1.097 - 5.484) | **0.0475** | 1.190 (0.4771 - 2.970) | 0.709 |
| TNM staging (II & III vs. I) | 2.71 (1.893 - 3.878) | **5.30E-08** | 2.647 (1.815 - 3.861) | **4.35E-07** |
| **Lung** |  |  |  |  |
| 8-gene signature (high vs. low risk) | 1.453 (1.036 - 2.038) | **0.0304** | 1.458 (1.041 - 2.043) | **0.0281** |
| TNM staging (II & III vs. I) | 1.597 (1.364 - 1.87) | **6.11E-09** | 1.598 (1.362 - 1.873) | **8.04E-09** |
| *TP53* mutation (Yes vs. No) | 1.316 (0.863 - 2.006) | 0.203 |  |  |
| *KRAS* mutation (Yes vs. No) | 0.764 (0.401 - 1.455) | 0.412 |  |  |
| *CDKN2A* mutation (Yes vs. No) | 1.759 (1.057 - 2.927) | **0.030** | 1.533 (0.920 - 2.554) | 0.101 |
| **Pancreas** |  |  |  |  |
| 8-gene signature (high vs. low risk) | 1.961 (1.212 - 3.174) | **0.00609** | 1.906 (1.177 - 3.089) | **0.00878** |
| TNM staging (II & III vs. I) | 1.339 (0.897 - 1.998) | 0.153 | 1.282 (0.835 - 1.967) | 0.255 |
| **Endometrium** |  |  |  |  |
| 8-gene signature (high vs. low risk) | 2.33 (1.341 - 4.05) | **0.0027** | 1.986 (1.136 - 3.472) | **0.016** |
| TNM staging (II & III vs. I) | 1.802 (1.431 - 2.269) | **5.60E-07** | 1.722 (1.365 - 2.172) | **4.56E-06** |

